# Chronic paternal morphine exposure increases sensitivity to morphine-derived antinociception

**DOI:** 10.1101/2021.02.07.430143

**Authors:** Andre B. Toussaint, William Foster, Jessica M. Jones, Samuel Kaufmann, Meghan Wachira, Robert Hughes, Angela R. Bongiovanni, Sydney T. Famularo, Benjamin P. Dunham, Ryan Schwark, Nathan T. Fried, Mathieu E. Wimmer, Ishmail Abdus-Saboor

## Abstract

Parental exposure to drugs of abuse such as opioids can have profound and long-lasting effects on reward processing and drug sensitivity across generations. However, little is known about the impact of long-term paternal exposure to morphine on offspring sensitivity to morphine-derived antinociception during painful experiences. To address this question, we constructed a rat pain scale at millisecond timescales to measure mechanical nociception in a multigenerational morphine exposure paradigm. Surprisingly, while developing the pain scale, we found that von Frey hair filaments (VFHs), the most common stimuli used in pain research, are not painful to rats and morphine did not change the touch-like responses elicited by VFHs. We next deployed this novel pain scale to determine whether chronic morphine exposure in sires impacted pain sensitivity in the next generation. Offspring produced from a cross of morphine-treated sires and drug-naïve dams, did not show any baseline changes in sensitivity to mechanical nociception. However, morphine-sired male progeny displayed a higher sensitivity to the antinociceptive properties of morphine, as measured by our pain scale. These findings demonstrate that long-term paternal exposure to morphine increases sensitivity to morphine-derived analgesia in the subsequent generation.

## Introduction

Growing evidence indicates that prenatal environmental factors can have profound and long-lasting influences on behavior and physiology in the next generations (Bale, 2015; Carone et al., 2010; Dias and Ressler, 2014; Goldberg and Gould, 2018; Kaletsky et al., 2020; Klengel et al., 2016; Shin Yim et al., 2017; Szutorisz et al., 2014; Yohn et al., 2015). Paternal opioid exposure is known to alter memory, reward processing, anxiety, and aggression in offspring (Ellis et al., 2020; Farah Naquiah et al., 2016; Goldberg and Gould, 2019; Vassoler et al., 2020). However, relatively little is known about the impact on pain sensitivity or antinociception in generations produced after chronic paternal morphine exposure. This gap in knowledge has major clinical implications in light of the current opioid epidemic – which is driven by unmet needs in resolving chronic pain (Hauser et al., 2017; Volkow and McLellan, 2016). It is possible that an individual’s usage of opiates like morphine, is partially influenced by their father’s exposure to these drugs, even before their birth. Longitudinal studies that directly probe this possibility are extremely difficult to pursue in clinical populations, largely because of their prohibitively high cost. Preclinical approaches have proven to be effective in systematically defining the phenotypes resulting from chronic opioid exposure and allow for assessment of the biological underpinnings of epigenetic inheritance of environmental insults. However, the question of morphine sensitivity to analgesia across generations remains unresolved, in part, because the tools commonly used to measure mechanical nociception in rats provide rather limited resolution.

A prerequisite for mechanistic pain discoveries in rodents are unbiased, quantitative, and reliable pain measurements that are consistent across time and research programs. Over the past fifty years, the field of pain research has relied primarily upon reflexive paw withdrawal assays with paw withdrawal frequency, latency, and threshold serving as a behavioral proxy for pain sensitivity (Le Bars et al., 2001; Mogil, 2009). In these studies, binary responses in mice or rats are the primary endpoints utilized to measure pain. However, rodents show paw withdrawal responses to both innocuous and noxious stimuli, highlighting the limitations of relying solely on paw withdrawal rates (Bourane et al., 2015; Duan et al., 2014; Murthy et al., 2018; Ranade et al., 2014). Other approaches to measure pain, such as operant conditioning or grimace scales may be better suited to define spontaneous pain and the negative affect associated with pain. However, these methods as currently constructed, are time consuming and relatively difficult to establish. For example, several elegant recent studies required a long training period (4-14 days) for semi-restrained and water deprived mice to self-report a sensory stimulus to the paw with a water reward, as a top-down indicator of the animal’s sensory state (Neubarth et al., 2020; Paricio-Montesinos et al., 2020). The grimace scales for mice and rats that focus on eye and facial features offer advantages, but they are also labor-intensive and not conducive for high-throughput analyses across all rodents of differing coat colors (Langford et al., 2010; Rossi et al., 2020; Sotocinal et al., 2011). To address the question of mechanical nociception in rats in the context of multigenerational morphine exposure, an ideal platform would allow an experimenter to rapidly test unrestrained rodents and produce easily quantifiable readouts.

The evidence regarding paternal morphine exposure and testing subsequent morphine-derived antinociception in subsequent generations is sparse and not conclusive. For example, prior studies exposed parents of each sex to morphine, oral application of morphine followed by wash-out, or application of a single injection of high-dosage morphine in sires (Ashabi et al., 2018; Cicero et al., 1995; Eriksson et al., 1989; Pachenari et al., 2019). These studies do not separate paternal morphine exposure from potentially confounding maternal behaviors. Nor do these studies provide self-administered long-term morphine, which is more congruent with morphine use in the clinic for chronic pain treatment. Thus, existing studies may not be as translationally relevant and the single-metric mechanical nociception tools used too course for accurate pain measurement. The goal of this study was to leverage a translationally relevant model of multigenerational morphine exposure with a novel rat pain scale to provide a better understanding of this highly clinically relevant question.

Here, we used our expertise in constructing mouse pain scales (Abdus-Saboor et al., 2019; Jones et al., 2020), to develop an analogous scale in rats to address the question of multigenerational morphine-derived antinociception. We developed a rat pain scale using high-speed recording of rapid paw and eye kinematics at 2,000 frames per second (fps) when a rat withdraws its paw from application of eight distinct mechanical stimuli. We used both sexes of two of the most commonly used outbred rat strains: Sprague Dawley and Long Evans. While mapping sub-second behavioral responses to the innocuous and noxious mechanical stimuli, we focused on behavioral features related to the stimulated paw’s height and speed, the presence of paw shaking, guarding, jumping, and eye grimace. For dimensionality reduction, we used principal component analysis (PCA) to find a low-dimensional subspace that combined our individual behavioral features into an easily interpretable pain scale. Following the PCA approach, we used support vector machine learning to make predictions about the pain-like probabilities of a given trial, training the machine with the first principal component score of innocuous and noxious stimuli. In addition to touch and objectively painful pinpricks, we evaluated pain-like responses to von Frey hair filaments (VFH), the most commonly used stimuli in pain research. Lastly, we used our recently described multigenerational morphine exposure paradigm to measure pain and morphine-derived antinociception in the offspring of sires that chronically self-administered morphine (Ellis et al., 2020). Together, this work reveals that a quantitative rat pain scale that maps sub-second pain signatures is a robust assessment tool for measuring pain in rats, even across generations.

## Results

### A composite score of nocifensive behaviors reveals statistical differences between responses to innocuous and noxious stimuli

We used high-speed imaging to capture sub-second behavioral features of the rat paw withdrawal in response to a set of innocuous [cotton swab (CS) and dynamic brush (DB)] and noxious [light pinprick (LP) and heavy pinprick (HP)] natural mechanical stimuli that would distinguish touch versus pain sensation. Our goal was to identify behavioral signatures associated with the innocuous versus noxious stimuli, and use statistical and machine learning approaches to combine these features into a single index to build a graded pain scale designed for rats **(Figure 1A)**. Our previous work assessing *in vivo* dorsal root ganglion calcium imaging in mice confirmed that CS and DB preferentially triggered touch sensitive neurons, whereas LP and HP activated neurons that are sensitive to pain (Abdus-Saboor et al., 2019). To capture these nuanced ethograms, the four stimuli were applied to the plantar surface of a randomly chosen hind-paw of fully acclimated adult Long Evans and Sprague Dawley rats of both sexes (40 rats in total; 10 for each group). To avoid sensitization, stimulus administration occurred over four days, with each day dedicated to one stimulus. High-speed imaging at 2,000 frames per second (fps) recorded withdrawal as well as whole-body movement, with a focus primarily on the stimulated hind-paw. High-speed videos captured nocifensive facial and behavioral features that are known to indicate pain sensation and follow a time sequence that lasted several seconds. In a typical example time series (**Figure 1B**), after administration of a noxious stimulus, the initial response of the rat involved an eye grimace or orbital tightening of its eyelids (captured at ~42 milliseconds), which is consistent with previously reported painful responses in rodents(Langford et al., 2010). Next, the rat would raise its hind paw away from the stimulus, hold it at the apex, then vigorously exhibit a sinusoidal paw shake (captured at ~ 70ms). Some rats also jumped in the air immediately after being stimulated (at ~100ms). After landing, the rat would then orient its head towards the pain-evoking stimulus in order to guard the affected paw (**Figure 1B,** captured at 621ms) (see **supplemental video 1** for example of the four nocifensive behaviors scored in high-speed). These four individual behavioral features - eye grimace/orbital tightening, paw shake, jumping, and paw guarding - were combined into one composite nocifensive score for each rat sex and strain.

**Figure 1.**
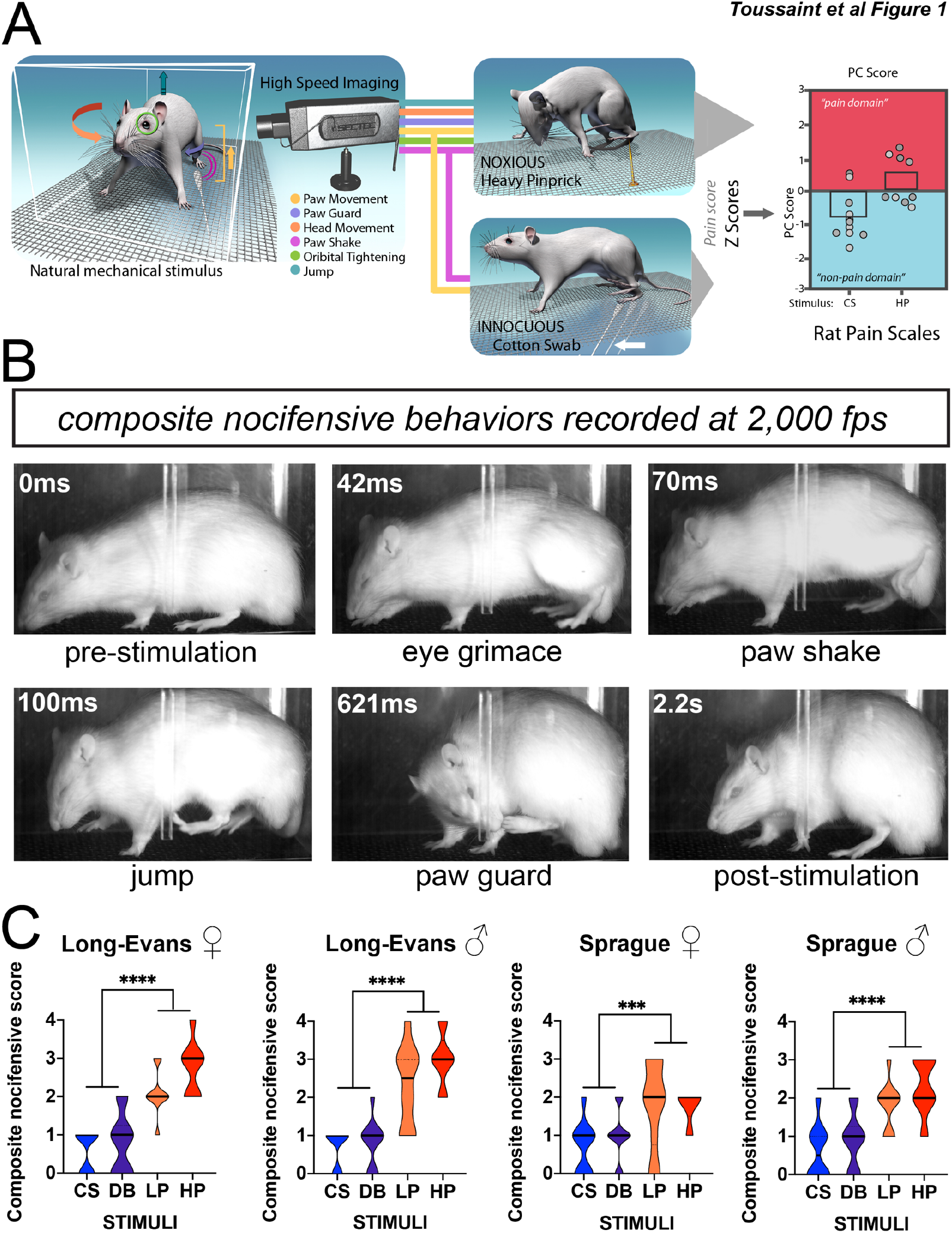
Temporal mapping of rat behavioral profile in response to mechanical stimuli. (A) Schematic of behavioral procedure and analysis showing lateral placement of high-speed camera in relation to freely behaving rat. Note that the camera lens captures the entire lateral area of the rat. (B) Representative single-frame images taken from high-speed videos of a Sprague Dawley rat following stimulation with a light pinprick. Each frame captures a distinct nocifensive behavior: eye grimace/orbital tightening (42ms), paw shake (70ms), jumping (100ms), paw guard (621ms). (C) Composite nocifensive score of Long Evans and Sprague Dawley female and male rats (40 rats in total; 10 per group) following stimulation with either a cotton swab (CS), dynamic brush (DB), light pinprick (LP), or heavy pinprick (HP). The score is a composite measurement of eye grimace/orbital tightening, paw shake, jumping, and paw guard. Truncated violin plots show the distribution density of the composite scores, with the solid horizontal top and bottom lines representing the minimum and maximum values, respectively; dashed horizontal line showing the median value; and dotted top and bottom horizontal lines representing the cutoff for the top 75% and bottom 25% percentile of values, respectively. *** p< 0.0001 and ** p< 0.05.

One-way ANOVA results revealed a marked difference in nocifensive responses to innocuous and noxious stimuli. Hind paw stimulation with CS or DB produced significantly lower behavioral scores than LP or HP across both sex and strain (**Figure 1C**; Long Evans Female: *F*(2.103, 23.13) = 22.06, p<0.0001; Long Evans Male: *F*(1.947, 16.88) = 27.37, p<0.0001; Sprague Dawley Female: *F*(1.724, 20.11) = 4.969, p=0.0213; Sprague Dawley Male: *F*(2.208, 17.67) = 14.47, p<0.0001). Interestingly, when stimulated with HP, Long Evans male rats displayed higher composite nocifensive scores than Sprague female and male rats that were stimulated with either LP or HP (**Figure 1C;** Long Evans Males HP vs. Sprague Dawley Females LP: p= 0.0342; Long Evans Males HP vs. Sprague Dawley Females HP: p= 0.0365; Long Evans Males HP vs. Sprague Dawley Males LP: p= 0.0342). These results support previous findings that suggest Long Evans rats are hypersensitive to mechanical stimuli (Mills et al., 2001). Taken together, this high-speed video imaging allowed us to capture subtle affective components that distinguish behaviors in response to touch and painful stimuli in Long Evans and Sprague Dawley rats.

### Rapid paw-kinematics demonstrate differences in responses to innocuous and noxious stimuli

To further quantify rat pain behavior, we next measured a set of parameters related to paw kinematics in response to innocuous and noxious stimuli **(Figure 2A)**. Based on our earlier work in mice (Abdus-Saboor et al., 2019; Jones et al., 2020), we anticipated that the paw lift speed and maximum paw height of the first lift would be sufficient to quantify kinematic movements for each stimulus trial that could inform the animal’s internal pain state. Following mechanical stimulation to the hind paw, we extracted the paw speed, by quantifying the distance from the initial paw lift to the highest point then dividing that value by the time in seconds between the two points.

**Figure 2.**
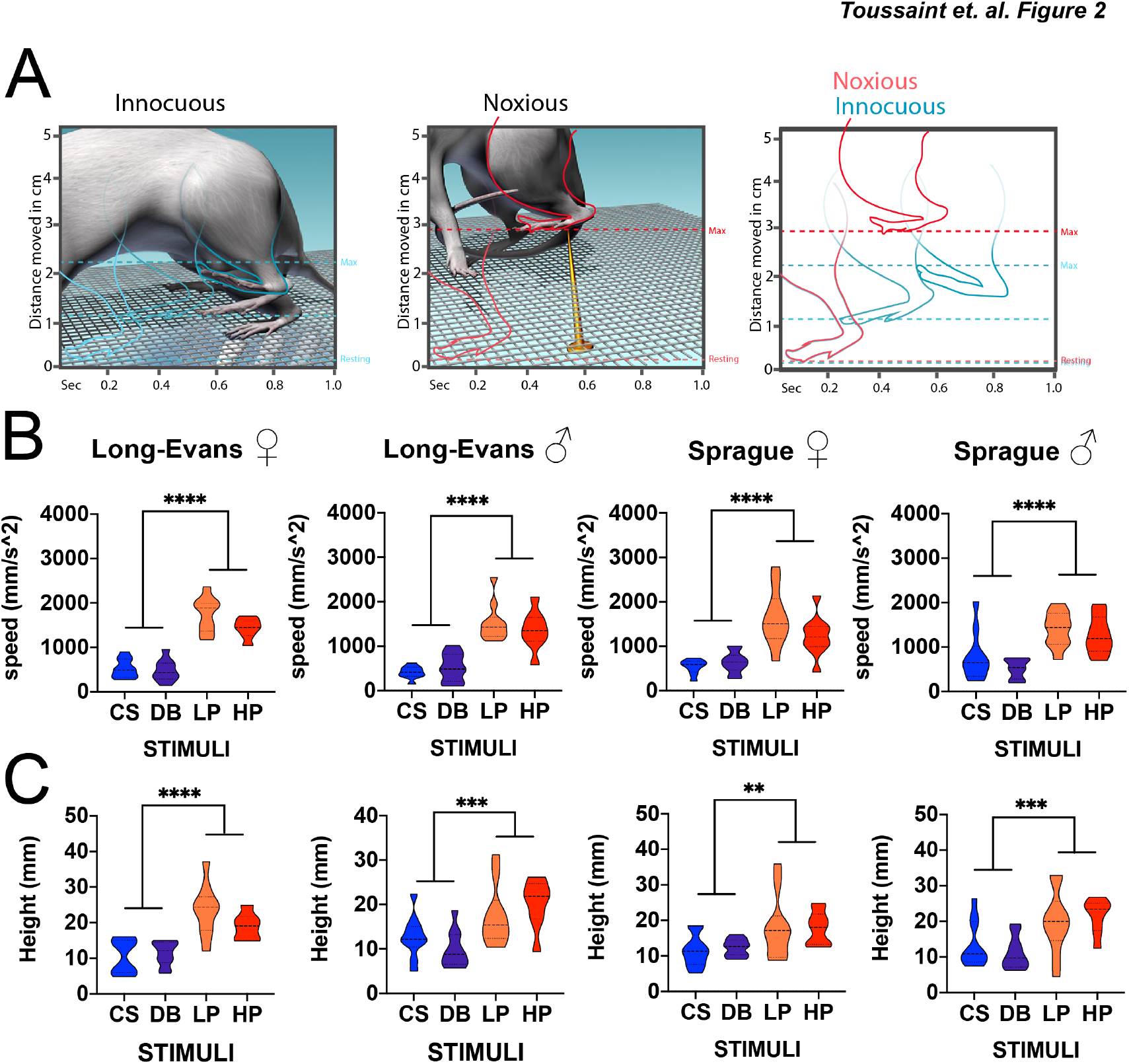
Sub-second temporal mapping of rat paw kinematics in response to mechanical stimuli. (A) Schematic of hind paw kinematic movements evoked by innocuous (CS and DB) or noxious (LP and HP) stimuli. Paw height (shown on the y-axis) is the distance from the mesh floor to the highest point following paw stimulation, while paw speed (shown on the x-axis) is the distance from the initial paw lift to the highest point divided by the time in seconds between the two points. (B) Paw speed of the first paw raise of the stimulated paw in Long Evans and Sprague Dawley female and male rats. (C) The maximum paw height of the first paw raise of the stimulated paw in Long Evans and Sprague Dawley female and male rats. Truncated violin plot shows the distribution density of the composite scores, with the solid horizontal top and bottom lines representing the minimum and maximum values, respectively; dashed horizontal line for the median value; and dotted top and bottom horizontal lines representing the cutoff for the top 75% and bottom 25% percentile of values, respectively. **** p< 0.0001, *** p< 0.001 and ** p< 0.05.

One-way ANOVAs revealed that the lift speed of the rat’s paw was significantly higher for noxious stimuli (LP, HP) compared to control innocuous stimuli (CS, DB) in female and male rats of both strains (**Figure 2B;** Long Evans Female: *F*(2.223, 23.71) = 48.24, p<0.0001; Long Evans Male: *F*(1.755, 19.89) = 25.84, p<0.0001; Sprague Dawley Female: *F*(1.995, 17.29) = 15.04, p=0.0002; Sprague Dawley Male: *F*(1.611, 18.26) = 9.737, p=0.0022). Although latency to withdraw their paw increased with stimulus intensity, no statistical difference was revealed for sex or strain in response to stimulation with a painful LP or HP (Sex and Strain: LP and HP: *F*(4.036, 36.90) = 1.454, p = 0.2358). These results suggest that both LP or HP evoked rapid withdrawal reflexes in Long Evans and Sprague Dawley rats of both sexes.

For the maximum paw height, we calculated the distance from the mesh floor to the highest point following hind paw stimulation. ANOVA analysis revealed that the behavioral kinematic responses evoked by a painful stimulus was greater than a non-painful stimulus for both sexes and strains examined (**Figure 2C;** Long Evans Female: *F*(1.958, 15.01) = 18.28, p<0.0001; Long Evans Male: *F*(1.866, 21.14) = 6.786, p=0.0061; Sprague Dawley Female: *F*(1.712, 14.84) = 3.922, p=0.0483; Sprague Dawley Male: *F*(2.070, 17.25) = 5.805, p=0.0112). Paw lift height increased in a stepwise manner with the application of each stimulus to the rat’s hind paw but no significant difference in sex or strain was found in response to application of the painful LP compared to HP (**Figure 2C;** Sex and Strain: LP and HP: *F*(2.927, 22.99) =0.8906, p=0.4586). Taken together, the millisecond paw kinematic responses to mechanical stimuli are sufficient to distinguish painful versus non-painful sensory states.

### Linear transformation and machine learning estimates the pain-probability of a given response on a trial-by-trial basis

Nocifensive behaviors and paw kinematics rely on unique and distinct features of the paw withdrawal response to consistently distinguish innocuous from painful stimuli. Each parameter holds a different dimension of statistical space, differing by unit and value of measurement. To comprehensively analyze and combine these measurements and to normalize each data point with respect to the population mean, we first converted all raw data to normalized z-scores **(Table 1)**. We subsequently combined all z-scores into a one-dimensional score using principle-component analysis (PCA), creating a PCA-generated pain score (PC1 score) that encompassed three behavioral dimensions and provided a threshold separating innocuous from noxious stimuli **(Figure 3A)**. We found that Long Evans and Sprague Dawley female and male rats had higher PCA-generated pain scores for the painful LP and HP compared to the innocuous CS and DB **(Figure 3B,** Long Evans Female: *F*(1.836, 14.07) = 63.14, p < 0.0001; Long Evans Male: *F*(2.155, 24.43) = 29.81, p < 0.0001; Sprague Dawley Female: *F*(1.544, 13.38) =10.20, p=0.0033; Sprague Dawley Male: *F*(2.001, 16.00) = 18.99, p<0.0001). Stated simply, positive PCA-generated pain scores indicate pain-like behavioral responses, while negative PCA-generated pain scores indicate innocuous behavioral responses. These scores provide a continuous gradation from most painful (high positive values) to most touch-like (high negative values) responses. Interestingly, in response to HP hind paw stimulation, the PCA-generated pain score for only Long Evans female rats were significantly higher than female Sprague Dawley rats, suggesting a strain difference in noxious stimuli for female rats (**Figure 3B,** Long Evans Females HP vs. Sprague Dawley Females HP: p=0.0146). Taken together, this PCA-generated scoring method in rats distinguished innocuous from noxious stimuli for a given trial, and it can be used to map a unique “pain state” in response to a painful pinprick in rats.

**Figure 3.**
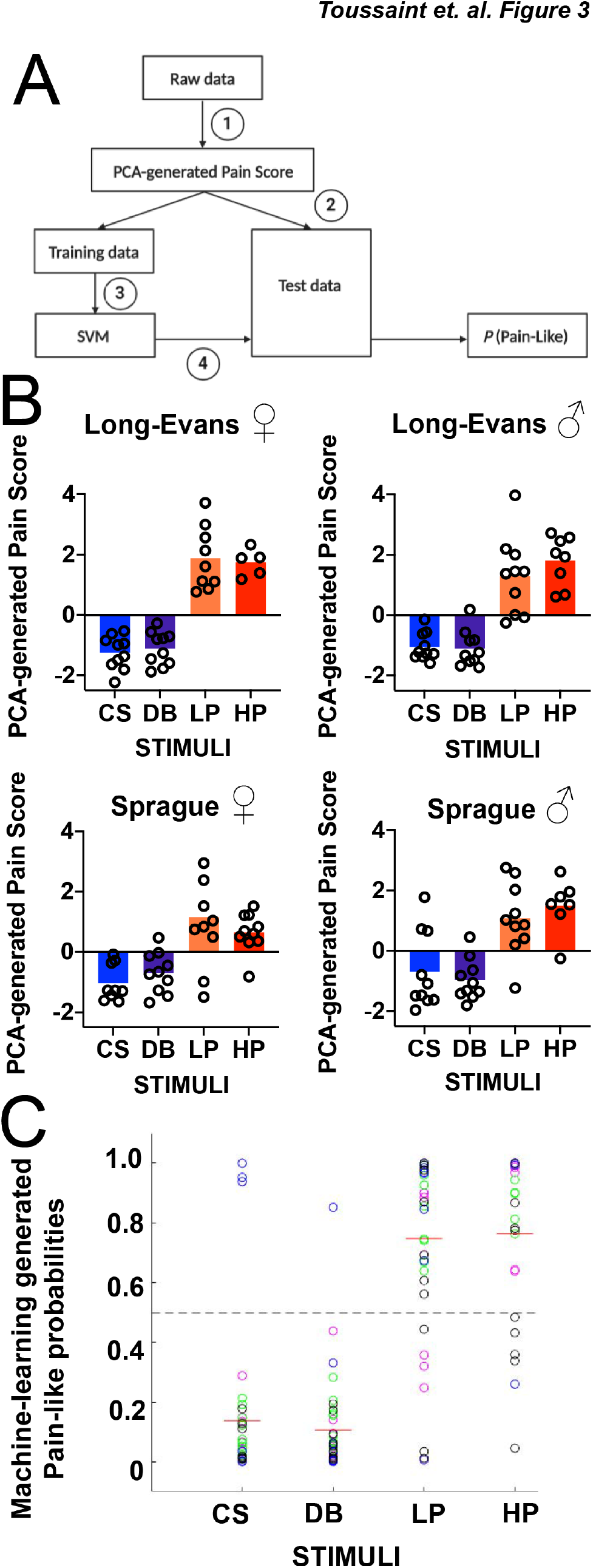
Trained support vector machine (SVM) predicts the pain-like probabilities for each stimulus. (A) SVM analytical pipeline. Step (1): calculate PC1 score of each trial by performing PCA on the z-scores from Table 1. Step (2): Split PC1 scores into training and testing sets. Step (3): train SVM with PC1 scores of training data (CS and HP). Step (4): predict pain-like probability (P [pain-like]) of remaining PC scores by training the model with three of the sex + strain combinations and testing on the remaining sex + strain. (B) PCA-generated pain scores for Long Evans and Sprague Dawley female and male rats following stimulation with innocuous (CS and DB) and noxious (LP and HP) stimuli. Negative values are indicative of “non-pain like” whereas positive values pain-like. (C) Machine learning-generated pain-like probabilities of CS, DB, LP, and HP trials in Long Evans and Sprague Dawley female and male rats. Color key: Green = Long Evans Females, Magenta = Long Evans Males, Black = Sprague Dawley Females, Blue = Sprague Dawley Males.

To further measure pain sensation based on behavioral responsivity, we used a machine-learning approach to predict the probability of each trial being pain-like. Principal Component 1 scores of CS and HP trials were used to train a support vector machine. We chose CS and HP trials because they triggered “non-painful” or “painful” behaviors with high confidence and the corresponding PCA scores showed the most consistent patterns across strain and sex. The trained SVM then predicted the probability of being “pain-like” for other trials **(Figure 3C)**. Thus, we could use machine learning to make predictions about unscored datasets. Quite analogously to the PCA based approach, we consistently separated innocuous versus noxious behavioral responses. Taken together, these statistical and machine learning approaches allow us to create easily interpretable pain scales - quantifying mechanical nociception in rats.

### Von Frey hair filaments do not elicit pain-like responses

Von Frey hair filaments (VFHs) are a frequently used stimulus to measure mechanical nociception in rodents, including in models of multigenerational opioid exposure. These stimuli are a mainstay in preclinical research, and yet discrepancies in the true nature of the sensation evoked by VFHs persists – evidenced by different labs considering the same force of VFH as painful or gentle (Francois et al., 2015; Woo et al., 2014). We used four VFHs ranging from the lowest to highest intensity traditionally used in the rat (**Figure 4A)**. Beginning with scoring the affective features that compose the nocifensive composite score (described in Figure 1), we did not observe any statistically significant difference across any of the VFHs (**Supplemental Figure 1A;** Long Evans Female: *F*(1.949, 17.54) = 2.168, p = 0.1450; Long Evans Male: *F*(2.324, 20.92) = 0.5000, p = 0.6408; Sprague Dawley Female: *F*(2.513, 22.61) =0.7663, p=0.5038; Sprague Dawley Male: *F*(2.435, 27.60) = 0.5330, p=0.6272). When assessing the kinematic movements of paw speed and height, only LE males showed a difference across VFH force for speed; all other comparisons revealed significant changes(**Supplemental Figure 1B,1C**; speed: Long Evans Female: *F*(1.529, 12.23) =0.6943, p=0.4807; Long Evans Male: *F*(1.905, 20.47) = 8.756, p=0.0020; Sprague Dawley Female: *F*(2.098, 24.48) = 3.029, p=0.0646; Sprague Dawley Male: *F*(2.189, 19.70) = 2.240, p=0.1294); height: Long Evans Female: *F*(2.294, 25.24) = 2.124, p = 0.1349; Long Evans Male: *F*(2.652, 30.06) =2.862, p=0.0592; Sprague Dawley Female: *F*(1.872, 21.84) = 1.690, p=0.2087; Sprague Dawley Male: *F*(2.110, 18.99) = 3.138, p=0.0642]).

**Figure 4.**
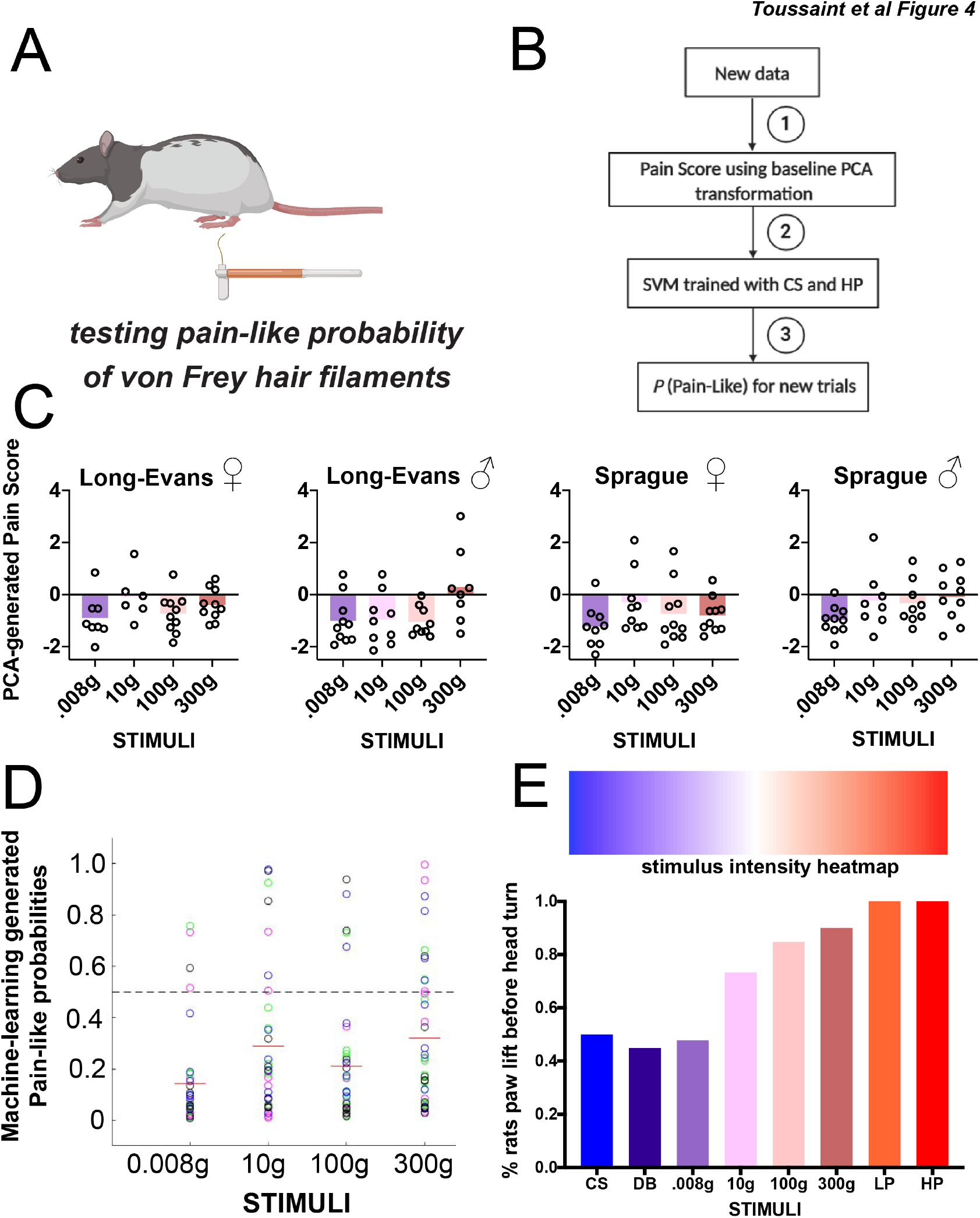
Sub-second temporal mapping of rat behavioral profile in response to von Frey hair filaments. (A,) Schematic and (B) Pain scale workflow: 1.) PC1 scores obtained using the previous baseline (CS,DB,LP,HP) PCA transformation. 2.) Predict on VF trials using the SVM trained on CS and HP baseline trials. 3.) generate probability pain-like on a trial by trial basis for the new data. (C) PCA-generated pain scores representing linear transformation of affective and reflexive behavioral features. Negative values are indicative of “non-pain like” whereas positive values pain-like. (D) Machine learning predictions made in Long Evans and Sprague Dawley female and male rats of pain-like probabilities of the same VFH filaments. Color key: Green = Long Evans Females, Magenta = Long Evans Males, Black = Sprague Dawley Females, Blue = Sprague Dawley Males. (E) Stimulus intensity heat map showing the percentage (y-axis) of rats that lifted their hind paw before turning to look at stimuli of increasing intensity. N=10 rats of each strain and sex per stimulus.

We next reduced the VF behavioral measurements into a single “pain scale” dimension using the data transformations derived from the PCA and machine learning approaches described above (**Figure 4B**). Corresponding to results obtained with the individual behavioral features (**Table 2**), the PCA-generated pain scores across sex and strain revealed mainly negative values, indicative of non-pain responses (**Figure 4C**). However, for males of each strain the PCA-generated pain scores were close to the threshold separating pain from non-pain, but not clearly painful like pinprick stimuli. The complementary machine-learning approach across strain and sex revealed that all VFHs have pain-like probabilities below 50%, indicating a low probability of being pain-like **(Figure 4D)**.

Given this unexpected result to stimulation with VFHs, we analyzed additional sub-second whole body features in animals that received these four VFHs, in addition to the four somatosensory stimuli described above (CS, DB, LP, HP). We focused on body orientation as a behavioral read-out, with the rationale that the body part to respond first to a stimulus may indicate how the animal is perceiving the stimulus. In other words, if an animal turns with its head to determine what has poked its paw before attempting to escape, the internal sensory state is likely non-painful. However, if an animal’s first response is to lift its paw in escape, before inquiring with head orientation to determine the origin of the stimulus, this is more likely reflecting an internal painful state. With this rationale, we interestingly saw a progressive increase in the proportion of animals that lifted their paws first that corresponded with stimulus intensity **(Figure 4E;** one way ANOVA: *F* (1.750, 5.251) = 6.728, p = 0.0375). We could align the stimuli in order from lowest to highest intensity based solely on this measurement (**Figure 4E**). Similar to our results with the PCA and machine learning approaches, the VFHs fell between the responses observed with dynamic brush and light pinprick. Taken together, these results demonstrate that at baseline conditions rats perceive VFHs more like touch stimuli, or just approaching the threshold separating touch from pain.

### Testing morphine-induced analgesia to VFHs across time using traditional scoring and the novel rat pain scale

The results from the previous set of experiments suggests that VFHs are not painful to rats at baseline conditions. If this is the case, how do we explain observations showing that morphine alters the threshold for rats to respond to VFHs? If one of the most well-known analgesics reduces VFH threshold sensitivity, is this sufficient evidence that VFHs must be painful to rats? Here, we decided to analyze these questions, using both traditional scoring and the new pain assessment tools described here in Long Evans male rats. This strain and sex was chosen because our current findings, as well as previous reports suggesting that these animals are more sensitive to mechanical stimulation (Mills et al., 2001). In this set of experiments, rats received baseline hind paw stimulation, followed by an immediate subcutaneous injection of morphine (3mg/kg). Subsequently, fifteen and sixty minutes post morphine injection, rats were tested again with the same stimuli(**Figure 5A)**.

**Figure 5.**
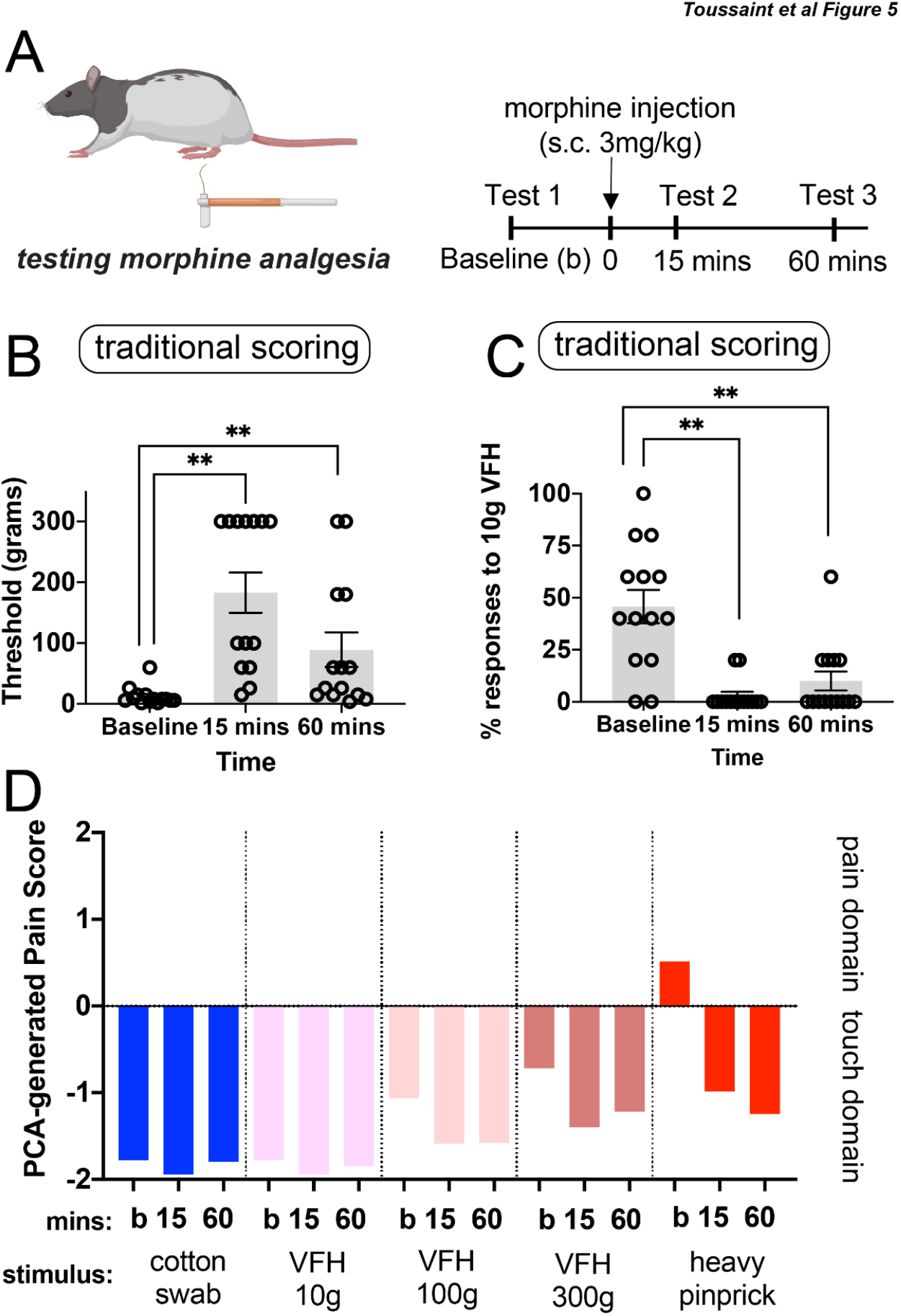
Using traditional scoring and our novel rat pain scale to measure morphine-induced analgesia over time in rats. (A) Schematic and Experimental timeline: Long Evans male rats received baseline (“b”) hind paw mechanical stimulations with VFHs or somatosensory stimuli. Rats then received a subcutaneous injection of morphine (3mg/kg) and were returned to their home cage. At 15-min and 60-min post morphine injection, rats were returned to the testing chamber to receive hind-paw stimulation using the same stimulus and procedure applied at baseline. (B) Traditional VFH thresholds after morphine treatment (3mg/kg). (C) Proportion of withdrawal responses at 10g VFH following morphine treatment. (D) Novel rat pain scale PCA-generated pain scores. * represents p< 0.05; ** represents p< 0.01 compared to baseline measurements

For the traditional scoring method, a series of VFHs were applied to the plantar surface of the hind paw in ascending order beginning with 1g force VFH. Threshold was defined as the force at which withdrawal responses occurred 40% of the time or more. If the threshold was below 10g, 5 stimulations were administered to define the percent response at 10g for each rat. Indeed, VFH threshold changed after morphine treatment (**Figure 5B,** Long Evans Males: *F*(1.934, 25.14) = 17.04, p < 0.0001). Sidak post-hoc analysis revealed an increase in threshold at 15 and 60 minutes post morphine injection compared to baseline (15 min, p= 0.0003 and 60 min, p=0.0281). Another way to analyze these data that is prominent in the field is to report the proportion of responses using the 10g VFH force, which has a baseline response rate of about 50% at baseline (Chaplan et al., 1994; Sandkuhler, 2009). Repeated measures ANOVA revealed that the proportion of withdrawal responses at 10g VFH stimulation decreased following morphine treatment (**Figure 5C,***F*(1.149, 14.94) = 16.12, p=0.0008). Sidak post hoc tests showed that the proportion of withdrawal responses was lower 15 minutes and 60 minutes after morphine treatment (3mg/kg) compared to baseline (15 minutes, p= 0.0006; 60 minutes p=0.0078). Taken together, these findings are consistent with the well characterized changes in response to VFH stimulations after morphine treatment (Bu et al., 2019; Mika et al., 2007; Popiolek-Barczyk et al., 2018; Williams et al., 2004).

We next used our novel rat pain scale to delineate the impact of morphine treatment on the response to VFH stimulations (**Table 2**). After morphine treatment, the PCA-generated pain scores at 15 and 60 minutes were unaffected (**Figure 5D;** F(1.949,25.22)=0.7187, p=0.4936). Stimulation with the 10g VFH as well as the 100g VFH elicited non pain-like PCA scores at baseline, and morphine treatment had no impact on their PCA-generated pain score (**Figure 5D**, 10g VFH: F(1.985,18.86)=0.2979, p=0.7442;100g VFH: F(1.721,24.09)=0.3466, p=0.0538). Interestingly, the 3mg/kg morphine treatment did produce a statistically significant effect with 300g VFH force (F(1.688,19.41)=3.931, p=0.0429), even though 300g mapped in the negative “non-pain” domain. Post-hoc Sidak tests revealed that the PCA scores in response to 300g VFH decreased 15 minutes after morphine treatment compared to baseline (p=0.0297). However there were no differences in PCA scores comparing 60 minute post morphine injection compared to baseline (p=0.2566). Using heavy pinprick as a known painful stimulus, we observed the initial hind paw stimulation generated a positive PCA score, consistent with a pain-like withdrawal reflex. Treatment with morphine had a significant analgesic response at 15 and 60 minutes after the injection (F(1.568,22.73)=20.33, p<0.001; post hoc sidak tests: 15 min: p=0.0027; 60 min: p<0.0001 compared to baseline). These experiments highlight the ability of the novel rat pain scale to detect morphine-derived antinociception using painful HP stimuli and reinforce our earlier observation that VFHs do not unambiguously register as painful stimuli to rats. Additionally, although morphine clearly alters how often a rat moves its paw to VFH stimulation, this does not automatically translate to a reduction in pain-like behaviors.

### Offspring of morphine-exposed sires show increased sensitivity to morphine-induced antinociception

With a new mechanical nociception measurement platform well validated, we next sought to address the long-standing question of whether chronic paternal morphine treatment altered morphine analgesia in offspring. Sires self-administered morphine for 60 days, while controls received saline. Sires were then bred to drug-naïve dams to produce first-generation (F1) offspring. Sires continued to self-administer morphine during the 5-day mating period to avoid withdrawal-related stress as a confounding factor. Pups were weaned at 21 days of age and group housed with same sex littermates until behavioral assessments as adults (60 + days of age, **Figure 6A,B)**. Baseline assessments using mechanical stimuli revealed no difference in pain-like responses in male morphine-sired animals compared to saline-sired controls (**Figure 6C-E**; Saline-sired males vs Morphine-sired males: CS: p=0.0748; DB: p=0.4929; VFH 10g: p=0.3095; VFH 100g: p=0.5480; VFH 300g: p=0.7157; LP: p=0.5145; HP: p=0.0862 (**Table 3**).

**Figure 6.**
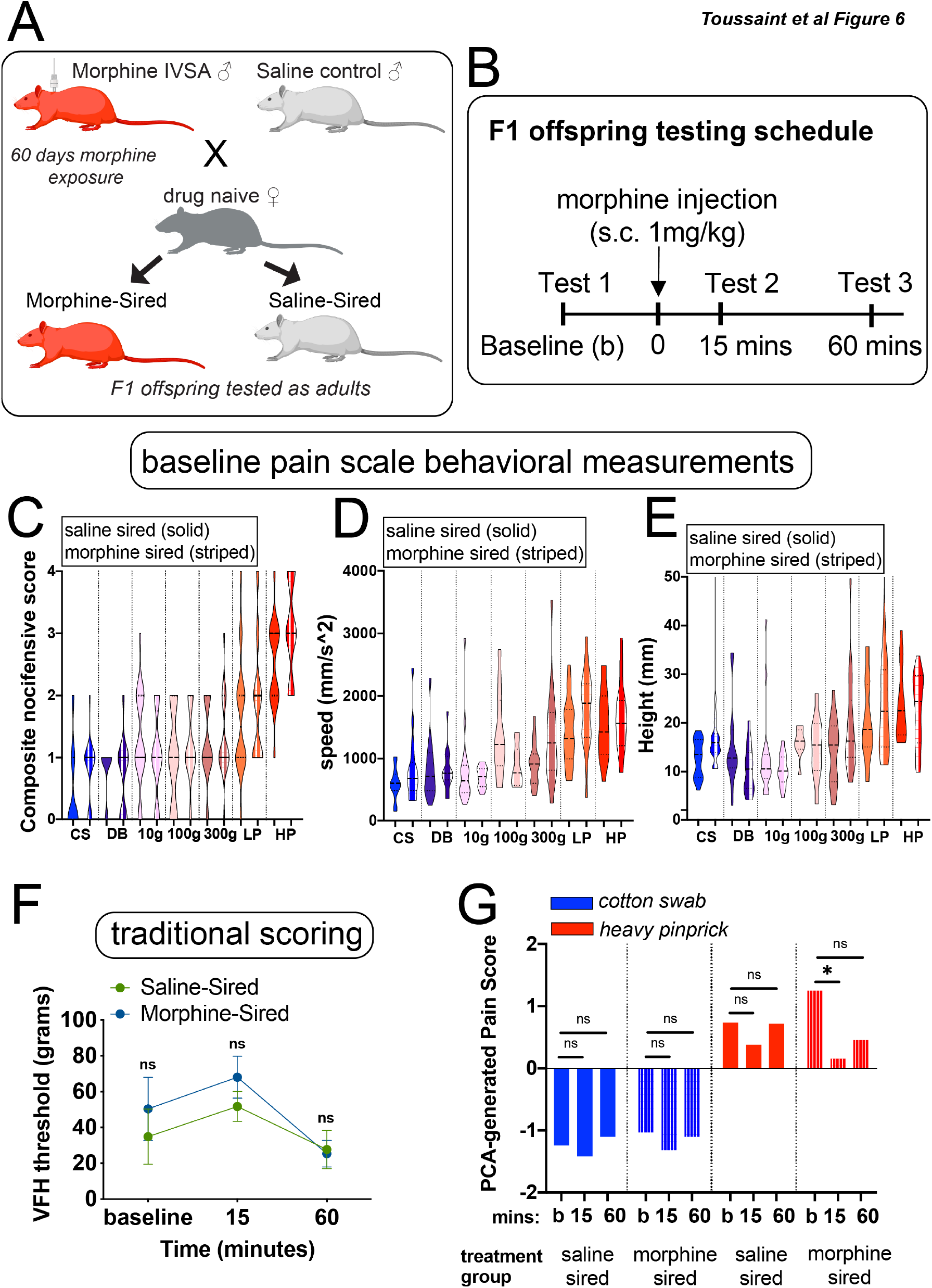
Multigenerational model of paternal morphine exposure used to assess pain in first-generation offspring. (A) Experimental paradigm and mating scheme for morphine intravenous self administration exposure. See methods for further details of the model. (B) Experimental timeline for injection protocol and behavioral testing. (C) Composite nocifensive score of F1 offspring following stimulation with the same baseline stimuli. (D,E) Hind paw speed and height kinematic movements evoked by the same stimuli. (F) Traditional scoring measuring VFH threshold does not separate treatment groups over time. (G) PCA-generated pain score predictions made in F1 offspring. Negative values are indicative of “non-pain like” whereas positive values pain-like. * represents p< 0.05.

We next examined whether the antinociceptive properties of morphine were affected by a paternal history of morphine exposure using both VFH-based traditional methods as well as our novel pain scale. Pain measurements were taken for male saline-sired and morphine-sired progeny and all animals were then injected with 1mg/kg morphine subcutaneously (**Figure 6B**). Pain measurements were also conducted 15 minutes and 60 minutes post morphine treatment. Using traditional methods, we found that the threshold for pain changed in both saline- and morphine-sired progeny after morphine treatment (**Figure 6F**, Sire: F(1,10)=0.61, p=.4529; time: F(2,20)=4.934, p=.0181; interaction: F(2,20)=0.4936, p=0.6177). Interestingly, post-hoc analyses revealed that neither saline-sired rats (p=0.4749, p=0.8691) nor morphine-sired rats (p=0.4420; p=0.2103) showed an increase in the VFH threshold 15 or 60 minutes after morphine injection compared to baseline. Importantly, these results suggest that paternal morphine exposure did not alter the sensitivity to morphine-derived antinociception. In the next experiment, we used a similar design combined with the novel pain scale. F1 male offspring received baseline mechanical stimulation to their hind-paw with 1 of 3 stimuli: cotton swab, light pinprick or heavy pinprick. Following this baseline recording, rats received a subcutaneous injection of low dose morphine (1mg/kg) and were returned to their home cage. Rats were tested again with the same stimulus 15 min and 60 min after the morphine injection. Behavioral responses were recorded using high-speed videography at all time-points. One-way ANOVA revealed that the nature of stimuli influenced the pain score regardless of siring (*F*(17, 150)=6.623, p < 0.0001), with pin pricks registering higher scores than innocuous stimuli. Morphine had no impact on the PCA-generated pain score in response to cotton swab stimulation (Time: F(1,26=1.34, p=0.2795; sire: F(1,18)=0.4932, p=0.4915; interaction: F(2,21)=0.4057, p=0.9603) for saline-sired (**Figure 6G**, 15 min: p=0.8220; 60 min p=0.8353 compared to baseline) or morphine-sired male progeny (15 min: p=0.5365; 60 min: p=0.9899 compared to baseline). For saline-sired offspring, this low dose of morphine did not significantly reduce the PCA-generated pain score in response to heavy pinprick (Time: F(1,28=5.149, p=0.0139; sire: F(1,15)=0.6263, p=0.4410; interaction: F(2,30)=1.272, p=0.2949; post-hoc-sidak tests: 15 min: p=0.2909; 60 min: p=0.9671 compared to baseline). In sharp contrast, morphine-sired male progeny displayed an antinociceptive response to 1mg/kg morphine after stimulation with l heavy pinprick (15 min: p=0.0370; 60 min: p=0.1039 compared to baseline). Interestingly, the composite nocifensive scores drove the PCA-generated pain scores, as the antinociceptive properties of morphine were most apparent in affective behaviors as compared to paw speed and height (**Supplemental Figure 2, Table 4**). Together, our pain scale demonstrated that the offspring of morphine-exposed sires are more sensitive to the antinociceptive properties of morphine – a finding that would have been missed with a traditional scoring method.

## Discussion

We have developed a rat pain scale recording sub-second mechanically evoked sensory behaviors at 2,000 fps with a focus on paw and eye movements following stimulus application. Recording and quantifying millisecond ethograms separated touch-like responses from pain-like responses in both sexes of two outbred rat strains. In addition, we transformed the multidimensional behavioral datasets into a single dimension with principal component analysis and machine learning to produce a comprehensive and quantitative rat pain scale. These approaches demonstrated that valence can be assigned to a response to a mechanical stimulus by focusing on the animal’s behavior, without making assumptions about how the animal might be experiencing the stimuli. Interestingly, the nature of the sensation evoked by stimulation with von Frey hair filaments was largely in the touch and not the painful domain as measured by this novel rat pain scale. This observation warrants further examination of the nature of VHFs and their utility in assessing pain at baseline conditions in rat models. Lastly, we used this novel analytical pipeline to assess the antinociceptive properties of morphine in a multigenerational model of paternal morphine exposure. Our findings demonstrate that the offspring produced by morphine-treated sires are hypersensitive to the antinociceptive effects of morphine compared to animals generated by sires exposed to saline.

### Behavior-centered approach allows statistical separation of touch versus pain in rats

Application of sensory stimuli to the plantar surface of the hind paw is the most common delivery system to measure mechanical pain in rats (Barrot, 2012). Withdrawal frequency or threshold are the predominant metrics for measuring the pain response when delivering mechanical stimuli to the rat paw. Our results here do not rely upon how many times the rat moves its paw, but rather we ask, “how does an animal respond behaviorally to the mechanical stimuli being delivered?” We find that behavioral responses to painful stimuli are driven by fast and high paw withdrawals, rapid eye grimacing, jumping, paw shaking and guarding, and removal of the paw prior to exploratory head movements. Thus, the simple paw withdrawal which is completed within seconds contains rich behavioral data that when mapped and quantified, allows statistical separation of responses that would appear identical to the unaided eye.

### von Frey hair filaments (VFHs) do not evoke pain in rats during baseline conditions

VFHs are the most common stimulus used in all of pain research, both in rodents, as well as in the clinic to test for peripheral sensory function (Barrot, 2012). When we constructed a mouse pain scale using an analogous approach to the work presented here, we noticed that the higher forces of VFHs (relative to mouse size), evoked clear painful behaviors (Abdus-Saboor et al., 2019). Therefore, it is possible that the responses to all stimuli may not be shared across species and assumptions about how rats will behave based on mouse studies should be treated with caution. For measuring mechanical allodynia following inflammatory or neuropathic pain, VFHs are probably still a useful mechanical stimulus because of their graded nature of delivering defined forces (Peirs et al., 2021; Wilson et al., 2019). However, when using VFHs during baseline conditions in rats, it is likely that researchers are measuring touch and not pain behavioral responses. What could account for the unexpected results obtained here about rats not perceiving VFHs as painful at baseline? While this remains unclear, it is possible that because the diameter of the filament at 300 grams is greatly expanded, the blunt nature of the stimulus would not be perceived as painful despite the heavy force. It is thus possible that 300 grams of force delivered by a more sharp filament, akin to the pinprick, would evoke a painful response to the rat.

Our results using the well-studied opiate morphine to blunt pain following application of somatosensory stimuli in rats also yielded unexpected results. Although morphine clearly reduced the paw withdrawal threshold to VFH stimuli, our pain scale determined that the behavioral responses to VFH were touch-like to begin with. How do we reconcile between these results? Our interpretation of these data is that at the dose of morphine injected, this had a slight sedative effect on the animal’s motor response without perturbing pain-like behaviors. This possibility is consistent with previous reports suggesting that morphine injections at these doses produce sedation in rats (Abbott and Guy, 1995; Fog, 1970; Kissin et al., 1990; Toyama et al., 2018). These findings demonstrate that it is possible to disentangle paw withdrawal from pain-like responses, and caution should be taken when attempting to link these two disparate measurements.

### Paternal morphine exposure has long-lasting consequences for morphine-derived antinociception in male progeny

Prenatal opioid exposure can have profound consequences for cognition, reward sensitivity and pain thresholds in the next generation (Vassoler and Wimmer, 2020). Relatively few multigenerational opioid exposure studies have focused on pain sensitivity in the next generation, but a consistent finding has been an increased sensitivity to opioid-induced antinociception in offspring produced by opioid-exposed sires and dams. Oral morphine administration in sires produced male offspring with increased sensitivity to morphine (7mg/kg) on a hot plate latency pain test (Eriksson et al., 1989). Male progeny derived from sires treated with a high dose of morphine (25mg/kg) acutely had increased sensitivity to the antinociceptive effect of opioids at higher doses (10 and 12mg/kg of morphine). In sharp contrast, female offspring produced by sires treated with morphine showed similar levels of morphine-derived analgesia (Cicero et al., 1995). Chronic morphine exposure of both dams and sires also increased the analgesic properties of a low dose (1.5mg/kg) of morphine in male progeny in a formalin-based pain assay (Ashabi et al., 2018). When paternal morphine exposure occurred during adolescence, the impact on nociception in male progeny was more subtle, mildly affecting baseline pain responses (Pachenari et al., 2018). Overall, it is clear that the nature and duration of parental or paternal morphine exposure is critical in determining the outcomes for the next generation.

Here, we focused on a highly translational model of opioid exposure using intravenous drug self-administration and an extended regimen that covers the sensitive period of spermatogenesis, which had never been done before. This approach offers a number of advantages, including the ability of sires to titrate the dose of morphine over this long duration and accounts for any development of tolerance over the course of the experiment. By narrowing our focus to the paternal lineage, we laid the groundwork for further investigation into the mechanisms underlying the transmission of the paternal morphine exposure. We have previously shown that maternal behavior is not impacted in dams bred to morphine-treated sires, which is a potential caveat in many of the aforementioned studies. Our unique approach combining the novel pain scale with a self-administration-based exposure protocol is an important first step in delineating whether and to what extent these observations may extend to human and clinical populations. Both the paternal exposure and the doses of morphine used to assess pain in progeny represent a substantial advance over previous research. Indeed these regimens are akin to doses used for pain management in patients. Moreover, the lower doses of morphine used in the current studies in the exposure model are not confounded by the potentially sedating effects associated with opioid treatments at higher doses (Abbott and Guy, 1995; Fog, 1970; Kissin et al., 1990; Toyama et al., 2018. Using traditional methods to measure pain, the differences in antinociception were missed in this multigenerational paradigm. Hence, the novel pain assessment pipeline outlined in this study was more sensitive to the effects of chronic paternal morphine exposure on morphine-derived anti-nociception in F1 progeny. This underscores the robustness of this new rapid approach to measuring mechanical nociception. Future studies will dive deeper into mechanistic detail, to uncover the identify of the molecule(s) that are transmitted from sperm to offspring, that increase sensitivity to morphine-derived antinociception.

## Methods and Materials

### Subject and Housing details

Sires and dams (F0 generation) were Sprague-Dawley and Long Evans rats weighing 250–300 g obtained from (Taconic Biosciences, Hudson, NY, USA). Sires were housed individually except for 1 week of pair housing during the mating period. Food and water were available ad libitum and rats were kept on a 12 h–12 h light–dark cycle. All experiments were performed during the light phase. All animal care and experiments were approved by the Institutional Animal Care and Use Committee of the University of Pennsylvania and Temple University and conducted following the National Institute of Health guidelines.

### Whole animal high speed videography

Rat behaviors were recorded at 2000 frames per second (fps) with a high-speed camera (Photron FastCAM Mini AX 50 170K-M-32GB - Monochrome 170K with 32GB memory) and attached lens (Zeiss 2/100M ZF.2-mount). Rats performed behavior in rectangular plexiglass chambers on an elevated mesh platform. The camera was placed at a ∼45° angle at ∼1-2 feet away from the Plexiglas holding chambers on a tripod with geared head for Photron AX 50. CMVision IP65 infrared lights that rats cannot detect were used to adequately illuminate the paw for subsequent scoring. Data collected on a Dell laptop computer with Photron FastCAM Analysis software.

### Mechanically evoked hind paw somatosensory behavioral paradigm

Consistent with previously published protocols, rats were acclimated to a rectangular plexiglas chamber where they could move freely but could not stand straight up. We delivered selected mechanical stimuli to the hind paw when rats were calm, still, and all four paws were in contact with the raised mesh platform. Rats habituated to the testing chambers before preforming behavioral tests. For the baseline experiments (CS, DB, LP, and HP) we tested the same number (10) of male and female Sprague Dawley and Long Evans rats, for a total of 40. We applied stimuli to the hind paw of each rat through the mesh floor. Cotton swab tests consisted of gentle contact between the cotton Q-tip and the hind paw of the mouse. Dynamic brush tests were performed by wiping a concealer makeup brush (e.l.f.™, purchased at the CVS) across the hind paw from back to front. We preformed light pinprick tests by gently touching a pin (Austerlitz Insect Pins®) to the hind paw of the mouse. We withdrew pins after achieving contact. We preformed heavy pinprick tests by sharply pressing this pin into the paw, pushing upward with force. The pin was withdrawn as soon as approximately 1/3 of the pins length had passed through the mesh. For application of von Frey hairs (VFHs, Stoelting Company, 58011), we used four different forces: 0.08 g, 10 g, 100 g, and 300 g. As previously described, we directed each VFH towards the center of the plantar paw and pressed upward until the filament bent (Cui et al., 2016). For the four natural stimuli and VFHs, a non-responsive animal did not respond within 2 s of stimulus delivery. For traditional measures using VFH, animals were placed in the same plexiglass enclosure and paw responses were scored as either withdrawn or not (binary outcome). Animals were stimulated 5 times with each of the following VFH: 1g, 2g, 4g, 6g, 8g, 10g, 15g, 26g, 60g, 100g, 180g, 300g. When animals withdrew their paw 2 or more times at a given force, the experiment stopped and the threshold was reported at that VFH force. If the threshold was below 10g, one more round of stimulation at 10g was performed to establish the percentage of responses at 10g. All experiments were carried out by an experimenter blind to experimental group.

### Scoring hind paw withdrawal movement features

We extracted aw height and paw speed from the high-speed videos and processed with Photron FastCAM software. We scored paw height in centimeters as the distance from the mesh floor to the highest point following paw stimulation. We calculated paw speed as the distance, in centimeters, from initial paw lift to the highest point, divided by the time in seconds between the two points. The composite nocifensive score is a composite of four individual behavior features: orbital tightening, paw shake, paw guard, and jumping. For example, if a given animal displayed 1/4 of those features it would receive a composite nocifensive score of 1. We scored orbital tightening when the eyes went from fully open to partially or fully closed following stimulus application. We defined paw shaking as high frequency paw flinching. We defined jumping as three or more paws off the mesh floor at the same time following a stimulus application. Lastly, we defined paw guard as any abnormal orientation, or placement, of the paw back during the descent of the withdrawal following stimulus application. We were not blind to the strain when scoring behaviors of wild-type rats, as Long Evans have white coats with a black hood color, while Sprague Dawleys are white-coated. However, we are blind to the stimulus type and VFH forces.

### Machine Learning

We classified paw withdrawal reflexes into “pain” and “non-pain” categories, using a principal component transformation of three measurements obtained from the high-speed imaging data: paw speed, paw height and composite noficensive score into a single dimension. A classification pipeline consisted of the following steps 1) the first principal component score (PC1) for each trial was derived from the z scores of all data, 2) the principal component 1 scores for the training data were used to train a support vector machine (SVM) with radial basis function kernels (kernel coefficient gamma = 1, penalty parameter C = 1), and 3) for a given trial, the SVM predicts the probability of that the response was pain-like based on that trial’s PC1 score. The data used to generate the PC1 scores and train the SVM for each figure can be seen in Table 1.

### Drugs

Morphine sulfate was a gift from the NIDA drug supply or obtained from Spectrum Chemical (Gardena, CA) and dissolved in sterile 0.9% saline.

### Multigenerational morphine exposure model

Adult male Sprague Dawley rats were anesthetized using an i.p. injection of a ketamine/xylazine cocktail (80 and 12 mg kg ^−1^, respectively). An indwelling silastic catheter (Instech Laboratories, Inc. Plymouth Meeting, PA) was fed into the right jugular vein, sutured in place, and mounted on the shoulder blade using a back-mount platform. Catheters were flushed daily with timentin (0.93 mg ml^−1^), dissolved in heparinized saline and sealed using metal caps, when not in use. Following catheterization surgery, rats were allowed to recover for 1 week then were placed in operant chambers to lever press for 3h daily access of infusions of morphine (0.75 mg/kg/infusion over 5 s) for sixty continuous days; control animals underwent the same catheterization surgeries and self-administration protocol but only had access to saline and were never exposed to morphine. Following chronic morphine self-administration, naïve female rats were placed in a cage with each male rat. Paternal stress has been shown to have long lasting consequences for offspring; therefore, sires continued to self-administer during the 5-day mating period to avoid withdrawal-related stress as a confounding factor (Morgan and Bale 2011, 2013, 2015). Sires were then removed and dams reared first-generation F1 progeny independently until post-natal day (PND) 21, at which point pups were weaned and group housed with littermates. Male and female offspring were pair-housed with same sex littermates upon weaning and remained pair-housed throughout behavioral testing. F1 behavioral testing was conducted when offspring were 2–6 months old. One to two animals from each litter were randomly selected for behavioral test, such that no litter was over-represented in any particular experiment. The Institutional Animal Care and Use Committee of Temple University approved all experiments.

### Timeseries experimental paradigm

Following 2-minutes of habituation inside of the testing chamber, naïve adult male rats received mechanical stimulation to their hindpaw for approximately 1-2 seconds or until an apparent behavioral response was elicited with 1 of 3 stimuli: cotton swab, light pinprick or heavy pinprick. A different stimulus was used for different groups of rats during the 3-day experiment, with the initial stimulation serving as a baseline measure of pain sensitivity. Rats were then immediately injected with morphine (1mg/kg, subcutaneously) and returned to their home cage. At 15-min and 60 min after the morphine injection, rats were returned to the testing chamber to receive hindpaw stimulation from the same stimulus. Behavioral responses were recorded using high-speed videography at baseline, 15min, and 60min post-injection.

### Quantification and Statistical Analysis

Following previously published protocols, we settled on three behavioral measures (paw height, paw speed, and composite nocifensive score). We performed dimension-reduction with a Principle Component Analysis using the paw height, paw speed, and composite nocifensive score. We could then combine normalized z-scores for each syllable into a single one-dimensional principle component for every stimulus trial.

---

**Supplemental Figure 1.**
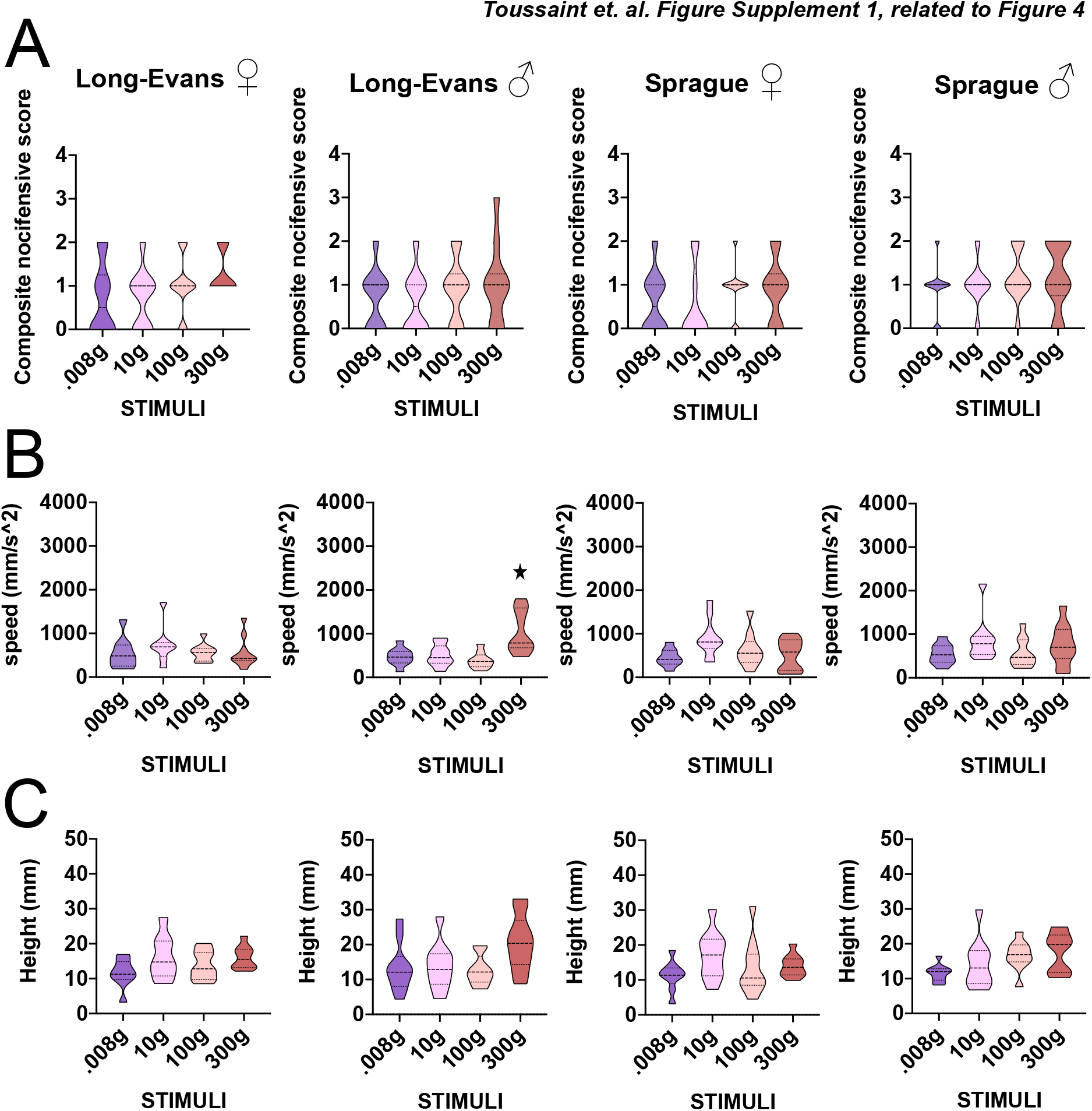
Sub-second temporal mapping of rat composite nocifensive score and paw kinematic behavioral profile in response to Von Frey hair filaments. (A) Composite nocifensive score of Long Evans and Sprague Dawley female and male rats (40 rats in total; 10 per group) following stimulation with Von Frey hair filaments of varying forces: 0.008, 10, 100, and 300 g. The score is a composite measurement of eye grimace/orbital tightening, paw shake, jumping, and paw guard. For instance, animals featuring 3 of the 4 behaviors are assigned a score of 3 for that particular trial. (B) Hind-paw kinematic movements evoked by the same VFH filaments. Paw speed of the first paw raise of the stimulated paw is the distance from the initial paw lift to the highest point divided by the time in seconds between the two points. (C) Paw height is the distance from the mesh floor to the highest point following paw stimulation. Truncated violin plot shows the distribution density of the composite scores, with the solid horizontal top and bottom lines representing the minimum and maximum values, respectively; dashed horizontal line for the median value; and dotted top and bottom horizontal lines representing the cutoff for the top 75% and bottom 25% percentile of values, respectively.

**Supplemental Figure 2.**
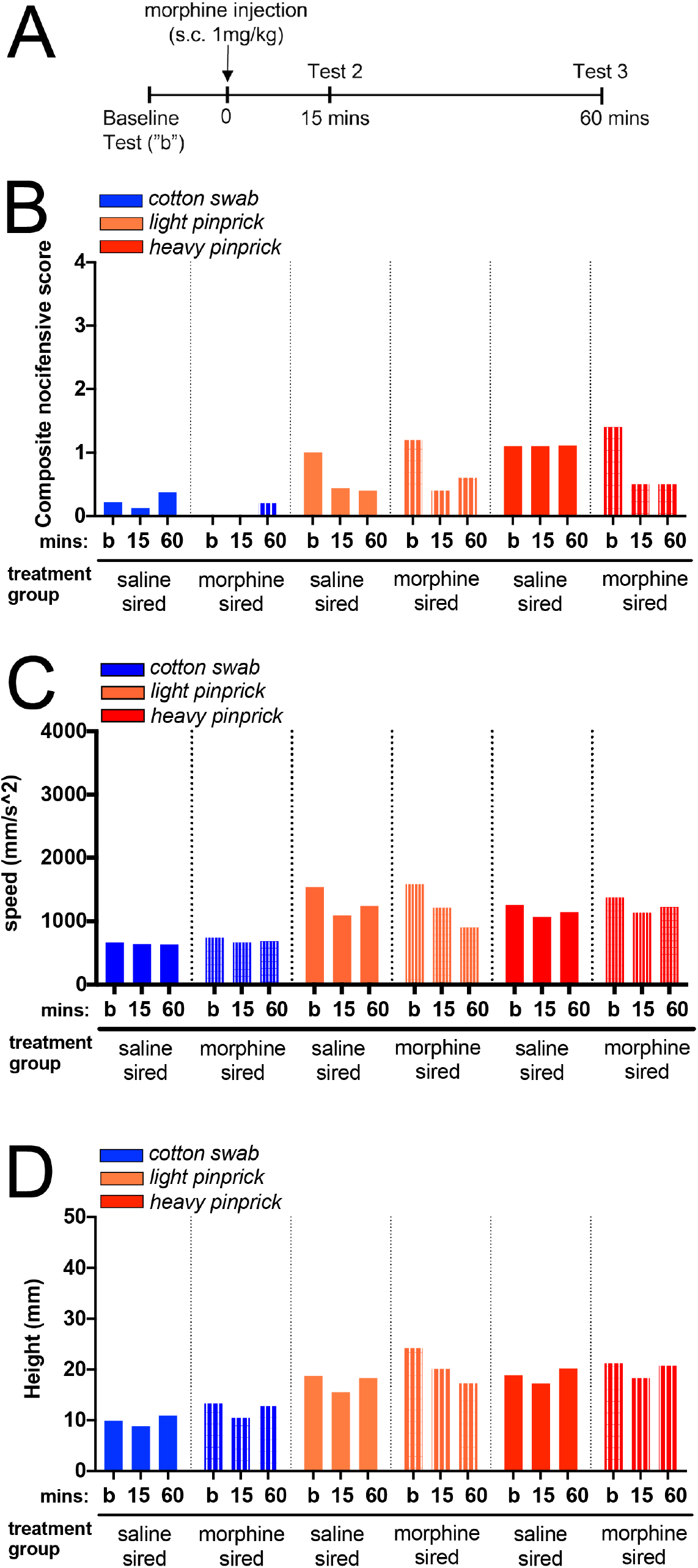
Baseline composite nocifensive score and paw kinematic behavioral profile of first-generation offspring derived from saline-exposed and morphine-exposed sires in response to application of mechanical stimuli to the hind paw. (A) Baseline composite nocifensive score drug-naïve first-generation male derived from sires exposed to wither saline or morphine. The score is a composite measurement of eye grimace/orbital tightening, paw shake, jumping, and paw guard. For instance, animals featuring 3 of the 4 behaviors are assigned a score of 3 for that particular trial. (B) Hind paw kinematic movements evoked by the same VFH filaments. Paw speed of the first paw raise of the stimulated paw is the distance from the initial paw lift to the highest point divided by the time in seconds between the two points. (C) Paw height is the distance from the mesh floor to the highest point following paw stimulation.

## Acknowledgements

We thank members of the Wimmer, Fried, and Abdus-Saboor labs for helpful discussion and comments on this manuscript. We thank Dr. Long Ding for consultation on the PCA and machine learning approaches adopted in this work. IA-S, JJ, and WF are supported by startup funds from the University of Pennsylvania and by a grant from the National Institutes of Health NIH/NIDCR, R00-DE026807, DP1 DA046537 (MEW), K01 DA039308 (MEW), T32 DA007237

(ABT; Unterwald EM, PI)

## Author contributions

N.F, M.W., and I.A. designed experiments, while all other authors performed experiments. All authors contributed to the analyses of the data and writing and editing of the manuscript.

